# Efficient and accurate causal inference with hidden con-founders from genome-transcriptome variation data

**DOI:** 10.1101/128496

**Authors:** Lingfei Wang, Tom Michoel

## Abstract

Mapping gene expression as a quantitative trait using whole genome-sequencing and transcriptome analysis allows to discover the functional consequences of genetic variation. We developed a novel method and ultra-fast software Findr for higly accurate causal inference between gene expression traits using cis-regulatory DNA variations as causal anchors, which improves current methods by taking into account hidden confounders and weak regulations. Findr outperformed existing methods on the DREAM5 Systems Genetics challenge and on the prediction of microRNA and transcription factor targets in human lymphoblastoid cells, while being nearly a million times faster. Findr is publicly available at https://github.com/lingfeiwang/findr.

**Author summary:** Understanding how genetic variation between individuals determines variation in observable traits or disease risk is one of the core aims of genetics. It is known that genetic variation often affects gene regulatory DNA elements and directly causes variation in expression of nearby genes. This effect in turn cascades down to other genes via the complex pathways and gene interaction networks that ultimately govern how cells operate in an ever changing environment. In theory, when genetic variation and gene expression levels are measured simultaneously in a large number of individuals, the causal effects of genes on each other can be inferred using statistical models similar to those used in randomized controlled trials. We developed a novel method and ultra-fast software Findr which, unlike existing methods, takes into account the complex but unknown network context when predicting causality between specific gene pairs. Findr’s predictions have a significantly higher overlap with known gene networks compared to existing methods, using both simulated and real data. Findr is also nearly a million times faster, and hence the only software in its class that can handle modern datasets where the expression levels of ten-thousands of genes are simultaneously measured in hundreds to thousands of individuals.

## 1 Introduction

Genetic variation in non-coding genomic regions, including at loci associated with complex traits and diseases identified by genome-wide association studies, predominantly plays a gene-regulatory role^1^. Whole genome and transcriptome analysis of natural populations has therefore become a common practice to understand how genetic variation leads to variation in phenotypes^2^. The number and size of studies mapping genome and transcriptome variation has surged in recent years due to the advent of high-throughput sequencing technologies, and ever more expansive catalogues of expression-associated DNA variants, termed expression quantitative trait loci (eQTLs), are being mapped in humans, model organisms, crops and other species^1,3–5^. Unravelling the causal hierarchies between DNA variants and their associated genes and phenotypes is now the key challenge to enable the discovery of novel molecular mechanisms, disease biomarkers or candidate drug targets from this type of data^6,7^.

It is believed that genetic variation can be used to infer the causal directions of regulation between coexpressed genes, based on the principle that genetic variation causes variation in nearby gene expression and acts as a causal anchor for identifying downstream genes^8,9^. Although numerous statistical models have been proposed for causal inference with genotype and gene expression data from matching samples^10–15^, no software implementation in the public domain is efficient enough to handle the volume of contemporary datasets, hindering any attempts to evaluate their performances. Moreover, existing statistical models rely on a conditional independence test which assumes that no hidden confounding factors affect the coexpression of causally related gene pairs. However gene regulatory networks are known to exhibit redundancy^16^ and are organized into higher order network motifs^17^, suggesting that confounding of causal relations by known or unknown common upstream regulators is the rule rather than the exception. Moreover, it is also known that the conditional independence test is susceptible to variations in relative measurement errors between genes^8,9^, an inherent feature of both microarray and RNA-seq based expression data^18^.

To investigate and address these issues, we developed Findr (Fast Inference of Networks from Directed Regulations), an ultra-fast software package that incorporates existing and novel statistical causal inference tests. The novel tests were designed to take into account the presence of unknown confounding effects, and were evaluated systematically against multiple existing methods using both simulated and real data.

## 2 Results

### 2.1 Findr incorporates existing and novel causal inference tests

Findr performs six likelihood ratio tests involving pairs of genes (or exons or transcripts) *A, B*, and an eQTL *E* of *A* (Table 1, Section 4.3). Findr then calculates Bayesian posterior probabilities of the hypothesis of interest being true based on the observed likelihood ratio test statistics (denoted *P_i_, i* = 0 to 5, 0 ≤ *P_i_* ≤ 1, Section 4.5). For this purpose, Findr utilizes newly derived analytical formulae for the null distributions of the likelihood ratios of the implemented tests (Section 4.4, **Figure S1**). This, together with efficient programming, resulted in a dramatic speedup compared to the standard computationally expensive approach of generating random permutations. The six posterior probabilities are then combined into the traditional causal inference test, our new causal inference test, and separately a correlation test that does not incorporate genotype information (Section 4.6). Each of these tests verifies whether the data arose from a specific subset of (*E, A, B*) relations (Table 1) among the full hypothesis space of all their possible interactions, and results in a probability of a causal interaction *A* → *B* being true, which can be used to rank predictions according to significance or to reconstruct directed networks of gene regulations by keeping all interactions exceeding a probability threshold.

**Table 1:**
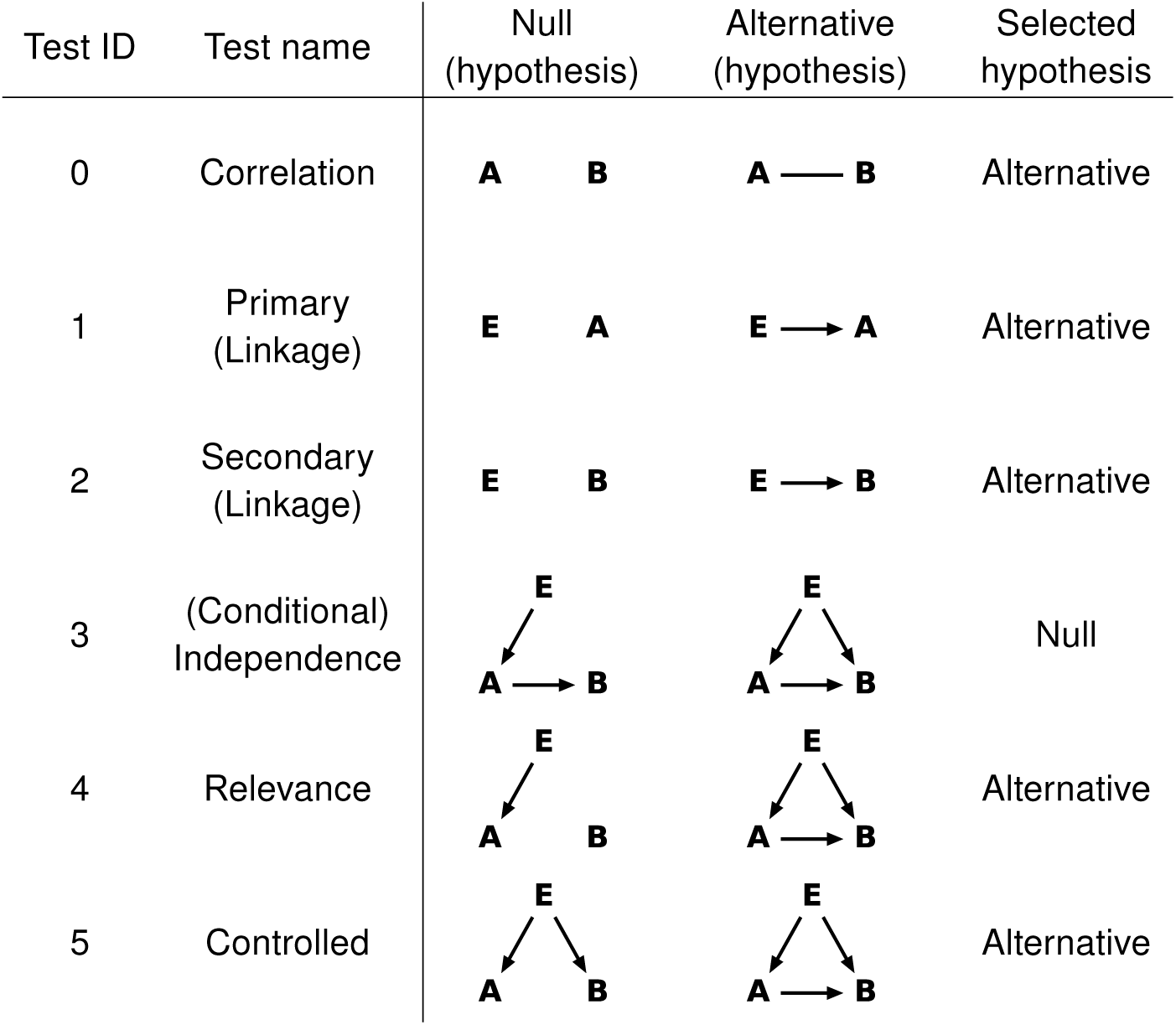
Six likelihood ratio tests are performed to test the regulation *A → B*, numbered, named, and defined as shown. *E* is the best eQTL of *A*. Arrows in a hypothesis indicate directed regulatory relations. Genes *A* and *B* each follow a normal distribution, whose mean depends additively on its regulator(s), as determined in the corresponding hypothesis. The dependency is categorical on discrete regulators (genotypes) and linear on continuous regulators (gene expression levels). The undirected line represents a multi-variate normal distribution between the relevant variables. In order to identify *A* → *B* regulation, we select either the null or the alternative hypothesis depending on the test, as shown.

### 2.2 Traditional causal inference fails in the presence of hidden confounders and weak regulations

Findr’s computational speed allowed us to systematically evaluate traditional causal inference methods for the first time. We obtained five datasets with 999 samples simulated from synthetic gene regulatory networks of 1,000 genes with known genetic architecture from the DREAM5 Systems Genetics Challenge (Section 4.1), and subsampled each dataset to observe how performance depends on sample size (Section 4.7). The correlation test (*P*_0_) does not incorporate genotype information and was used as a benchmark for performance evaluations in terms of areas under the receiver operating characteristic (AUROC) and precision-recall (AUPR) curves (Section 4.7). The traditional method^11^ combines the secondary (*P*_2_) and independence (*P*_3_) tests sequentially (Table 1, Section 4.6), and was evaluated by comparing *P*_2_ and *P*_2_*P*_3_ separately against the correlation test. Both the secondary test alone and the traditional causal inference test combination were found to *underperform* the correlation test (Figure 1A,B). Moreover, the inclusion of the conditional independence test *worsened* inference accuracy, more so with increasing sample size (Figure 1A,B) and increasing number of regulations per gene (**Supplementary Material S2.3**). Similar performance drops were also observed for the Causal Inference Test (CIT)^13,15^ software, which also is based on the conditional independence test (**Figure S3**).

**Figure 1:**
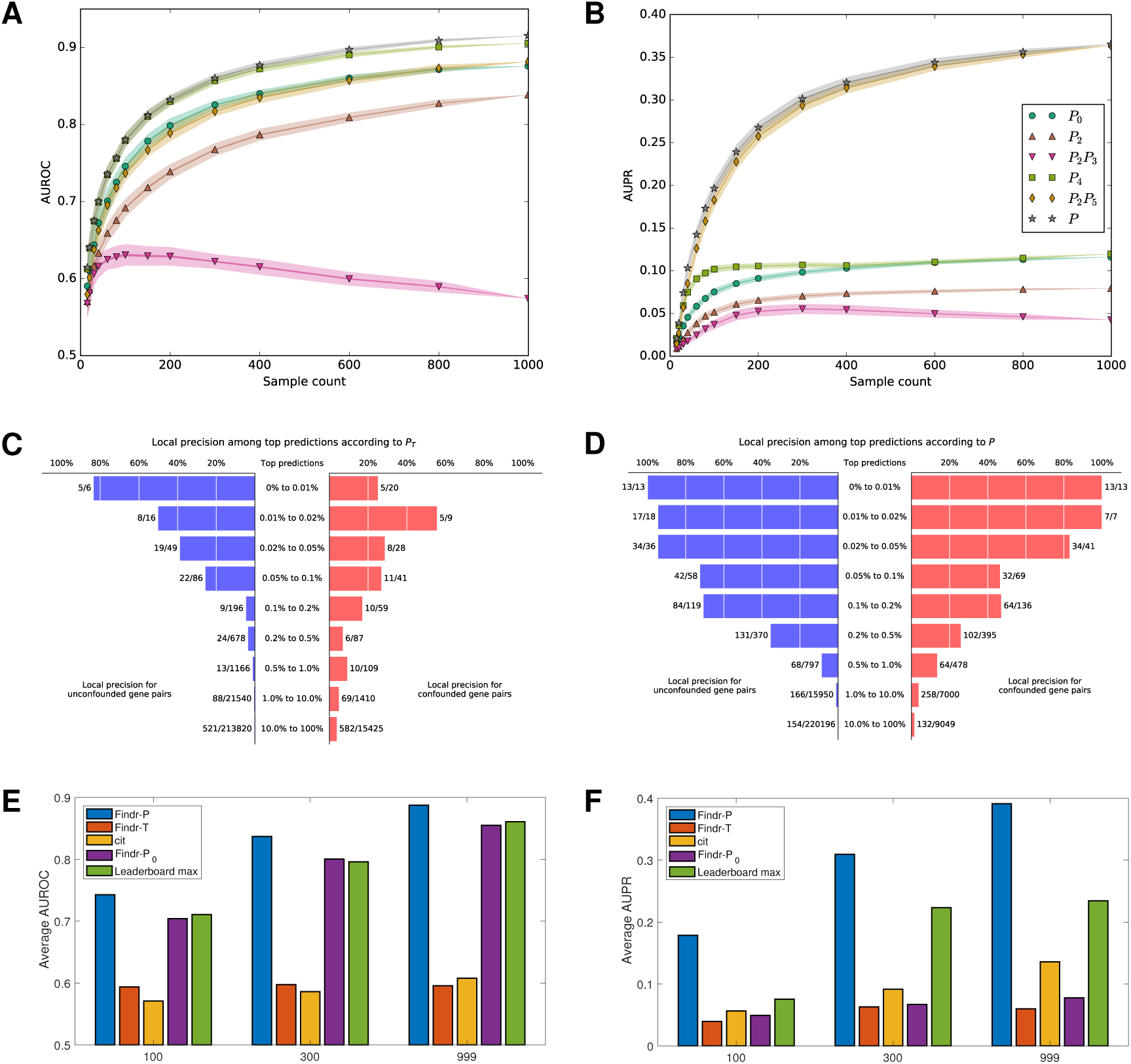
Findr achieves best prediction accuracy on the DREAM5 Systems Genetics Challenge. (**A**, **B**) The mean AUROC (**A**) and AUPR (**B**) on subsampled data are shown for traditional (*P*_2_, *P*_2_*P*_3_) and newly proposed (*P*_4_, *P*_2_*P*_5_, *P*) causal inference tests against the baseline correlation test (*P*_0_). Every marker corresponds to the average AUROC or AUPR at specific sample sizes. Random subsampling at every sample size was performed 100 times. Half widths of the lines and shades are the standard errors and standard deviations respectively. *P_i_* corresponds to test i numbered in Table 1; *P* is the new composite test (Section 4.6). This figure is for dataset 4 of the DREAM challenge. For results on other datasets of the same challenge, see **Figure S2**. (**C**, **D**) Local precision of top predictions for the traditional (**C**) and novel (**D**) tests for dataset 4 of the DREAM challenge. Numbers next to each bar (*x/y*) indicate the number of true regulations (*x*) and the total number of gene pairs (*y*) within the respective range of prediction scores. For results on other datasets, see **Figure S6**. (**E**, **F**) The average AUROC (**E**) and AUPR (**F**) over 5 DREAM datasets with respectively 100, 300 and 999 samples are shown for Findr’s new (Findr-*P*), traditional (Findr-*P*_*T*_), and correlation (Findr-*P*_0_) tests, for CIT and for the best scores on the DREAM challenge leaderboad. For individual results on all 15 datasets, see **Table S1**.

We believe that the failure of traditional causal inference is due to an elevated false negative rate (FNR) coming from two sources. First, the secondary test is less powerful in identifying weak interactions than the correlation test. In a true regulation *E* → *A* → *B*, the secondary linkage (*E* → *B*) is the result of two direct linkages chained together, and is harder to detect than either of them. The secondary test hence picks up fewer true regulations, and consequently has a higher FNR. Second, the conditional independence test is counter-productive in the presence of hidden confounders (i.e. common upstream regulators). In such cases, even if *E* → *A* → *B* is genuine, the conditional independence test will find *E* and *B* to be still correlated after conditioning on *A* due to a collider effect (**Figure S5**)^19^. Hence the conditional independence test only reports positive on *E* → *A* → *B* relations without confounder, further raising the FNR. This is supported by the observation of worsening performance with increasing sample size (where confounding effects become more distinguishable) and increasing number of regulations per gene (which leads to more confounding).

To further support this claim, we examined the inference precision among the top predictions from the traditional test, separately for gene pairs directly unconfounded or confounded by at least one gene (Section 4.7). Compared to unconfounded gene pairs, confounded ones resulted in significantly more false positives among the top predictions (Figure 1C). Furthermore, the vast majority of real interactions fell outside the top 1% of predictions (i.e. had small posterior probability) [92% (651/706) for confounded and 86% (609/709) for unconfounded interactions, Figure 1C]. Together, these results again showed the failure of the traditional test on confounded interactions and its high false negative rate overall.

### 2.3 Findr accounts for weak secondary linkage, allows for hidden confounders, and outperforms existing methods on simulated data

To overcome the limitations of traditional causal inference methods, Findr incorporates two additional tests (Table 1 and Section 4.3). The relevance test (*P*_4_) verifies that *B* is not independent from *A* and *E* simultaneously and is more sensitive for picking up weak secondary linkages than the secondary linkage test. The controlled test (*P*_5_) ensures that the correlation between *A* and *B* cannot be fully explained by *E*, i.e. excludes pleiotropy. The same subsampling analysis revealed that *P*_4_ performed best in terms of AUROC, and AUPR with small sample sizes, whilst the combination *P*_2_*P*_5_ achieved highest AUPR for larger sample sizes (Figure 1A,B). Most importantly, both tests consistently outperformed the correlation test (*P*_0_), particularly for AUPR. This demonstrates conclusively in a comparative setting that the inclusion of genotype data indeed can improve regulatory network inference. These observations are consistent across all five DREAM datasets (**Figure S2**).

We combined the advantages of *P*_4_ and *P*_2_*P*_5_ by averaging them in a composite test (*P*) (Section 4.6), which outperformed *P*_4_ and *P*_2_*P*_5_ at all sample sizes (Figure 1 and **Figure S2**) and hence was appointed as Findr’s new test for causal inference. Findr’s new test (*P*) obtained consistently higher levels of local precision (i.e. one minus local FDR) on confounded and unconfounded gene pairs compared to Findr’s traditional causal inference test (*P_T_*) (Figure 1C,D), and outperformed the traditional test (*P****_T_***), correlation test (*P*_0_), CIT, and every participating method of the DREAM5 Systems Genetics Challenge (Section 4.1) in terms of AUROC and AUPR on all 15 datasets (Figure 1E,F, **Table S1, Figure S4**).

Specifically, Findr’s new test was able to address the inflated FNR of the traditional method due to confounded interactions. It performed almost equally well on confounded and unconfounded gene pairs and, compared to the traditional test, significantly fewer real interactions fell outside the top 1% of predictions (55% vs. 92% for confounded and 45% vs. 86% for unconfounded interactions, Figure 1D, **Figure S6**).

### 2.4 The conditional independence test incurs false negatives for unconfounded regulations due to measurement error

The traditional causal inference method based on the conditional indepedence test results in false negatives for confounded interactions, whose effect was shown signficant for the simulated DREAM datasets. However, the traditional test surprisingly reported more confounded gene pairs than the new test in its top predictions (albeit with lower precision), and correspondingly fewer unconfounded gene pairs (Figure 1C,D, **Figure S6**).

We hypothesized that this inconsistency originated from yet another source of false negatives, where measurement error can confuse the conditional independence test. Measurement error in an upstream variable (called **A** in Table 1) does not affect the expression levels of its downstream targets, and hence a more realistic model for gene regulation is *E* → *A*^(t)^ → *B* with *A*^(t)^ → *A*, where the measured quantities are *E, A*, and *B*, but the true value for *A*, noted *A*^(t)^, remains unknown. When the measurement error (in *A*^(t)^ → *A*) is significant, conditioning on *A* instead of *A*^(t)^ cannot remove all the correlation between *E* and *B* and would therefore report false negatives for unconfounded interactions as well. This effect has been previously studied, for example in epidemiology as the “spurious appearance of odds-ratio heterogeneity”^20^.

We verified our hypothesis with a simple simulation (Section 4.8). In a typical scenario with 300 samples from a monoallelic species, minor allele frequency 0.1, and a third of the total variance of *B* coming from *A*^(t)^, the conditional independence test reported false negatives (likeilihood ratio p-value ≪ 1, i.e. rejecting the null hypothesis of conditional indepencence, cf. Table 1) as long as measurement error contributed more than half of *A*’s total unexplained variance (Figure 2B). False negatives occurred at even weaker measurement errors, when the sample sizes were larger or when stronger *A* → *B* regulations were assumed (**Figure S7**).

**Figure 2:**
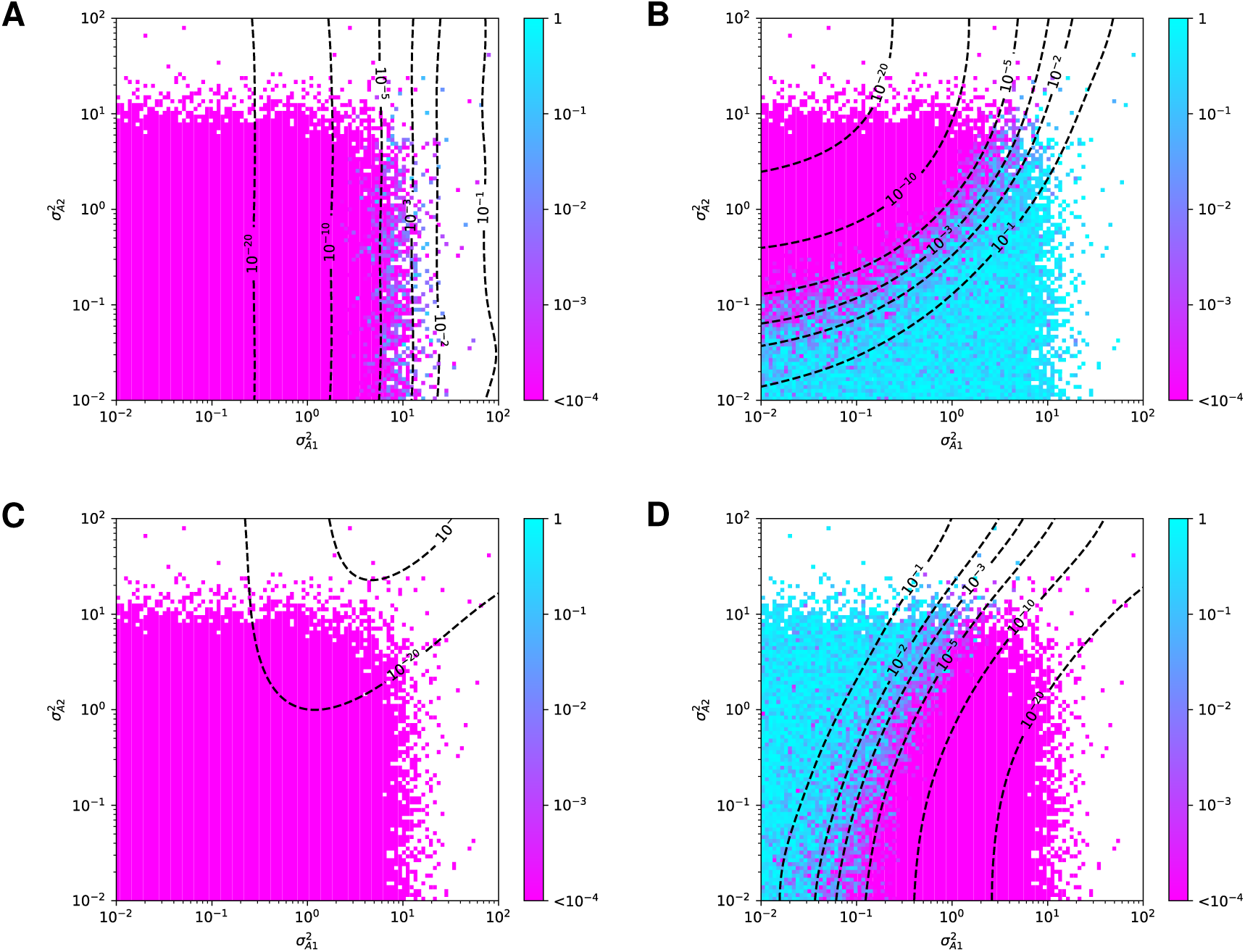
The conditional independence test yields false negatives for unconfounded regulations in the presence of even minor measurement errors. Null hypothesis p-values of the secondary linkage (**A**), conditional independence (**B**), relevance (**C**), and controlled (**D**) tests are shown on simulated data from the ground truth model *E* → *A*^(t)^ → *B* with *A*^(t)^ → *A*. *A*^(t)^’s variance coming from *E* is set to one, 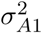 is *A*^(t)^’s variance from other sources and 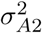 is the variance due to measurement noise. *A* total of 100 values from 10^−2^ to 10^2^ were taken for 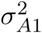 and 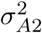 to form the 100 × 100 tiles. Tiles that did not produce a significant eQTL relation *E* → *A* with *p*-value ≤ 10^−6^ were discarded. Contour lines are for the log-average of smoothened tile values. Note that for the conditional independence test (**B**), the true model corresponds to the null hypothesis, i.e. small (purple) p-values correspond to *false negatives*, whereas for the other tests the true model corresponds to the alternative hypothesis, i.e. small (purple) p-values correspond to *true positives* (cf. Table 1). For details of the simulation and results from other parameter settings, see Section 4.8 and **Figure S7** respectively.

This observation goes beyond the well-known problems that arise from a large measurement error in all variables, which acts like a hidden confounder^9^, or from a much larger measurement error in *A* than *B*, which can result in *B* becoming a better measurement of *A*^(t)^ than *A* itself^8^. In this simulation, the false negatives persisted even if *E* → *A* was observationally much stronger than *E* → *B*, such as when *A*’s measurement error was only 10% (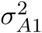 = 0.1) compared to up to **67%** for *B* (Figure 2B). This suggested a unique and mostly neglected source of false negatives that would not affect other tests. Indeed, the secondary, relevance, and controlled tests were much less sensitive to measurement errors (Figure 2A,C,D).

### 2.5 Findr outperforms traditional causal inference and machine learning methods on microRNA target prediction

In order to evaluate Findr on a real dataset, we performed causal inference on miRNA and mRNA sequencing data in lymphoblastoid cell lines from 360 European individuals in the Geuvadis study^3^ (Section 4.1). We first tested 55 miRNAs with reported significant cis-eQTLs against 23,722 genes. Since miRNA target predictions from sequence complimentarity alone result in high numbers of false positives, prediction methods based on correlating miRNA and gene expression profiles are of great interest^21^. Although miRNA target prediction using causal inference from genotype and gene expression data has been considered^22^, it remains unknown whether the inclusion of genotype data improves existing expression-based methods. To compare Findr against the state-of-the-art for expression-based miRNA target prediction, we used miRLAB, an integrated database of experimentally confirmed human miRNA target genes with a uniform interface to predict targets using twelve methods, including linear and non-linear, pairwise correlation and multivariate regression methods^23^. We were able to infer miRNA targets with 11/12 miRLAB methods, and also applied the GENIE3 random forest regression method^24^, CIT, and the three tests in Findr: the new (*P*) and traditional (*P*_*T*_) causal inference tests and the correlation test (*P*_0_) (**Supplementary Material S2.4**). Findr’s new test achieved highest AUROC and AUPR among the 16 methods attempted. In particular, Findr’s new test significantly outperformed the traditional test and CIT, the two other genotype-assisted methods, while also being over 500,000 times faster than CIT (Figure 3, **Table S2**, **Figure S8**). Findr’s correlation test outperformed all other methods not using genotype information, including correlation, regression, and random forest methods, and was 500 to 100,000 times faster (Figure 3, **Table S2**, **Figure S8**). This further illustrates the power of the Bayesian gene-specific background estimation method implemented in all Findr’s tests (Section 4.5).

**Figure 3:**
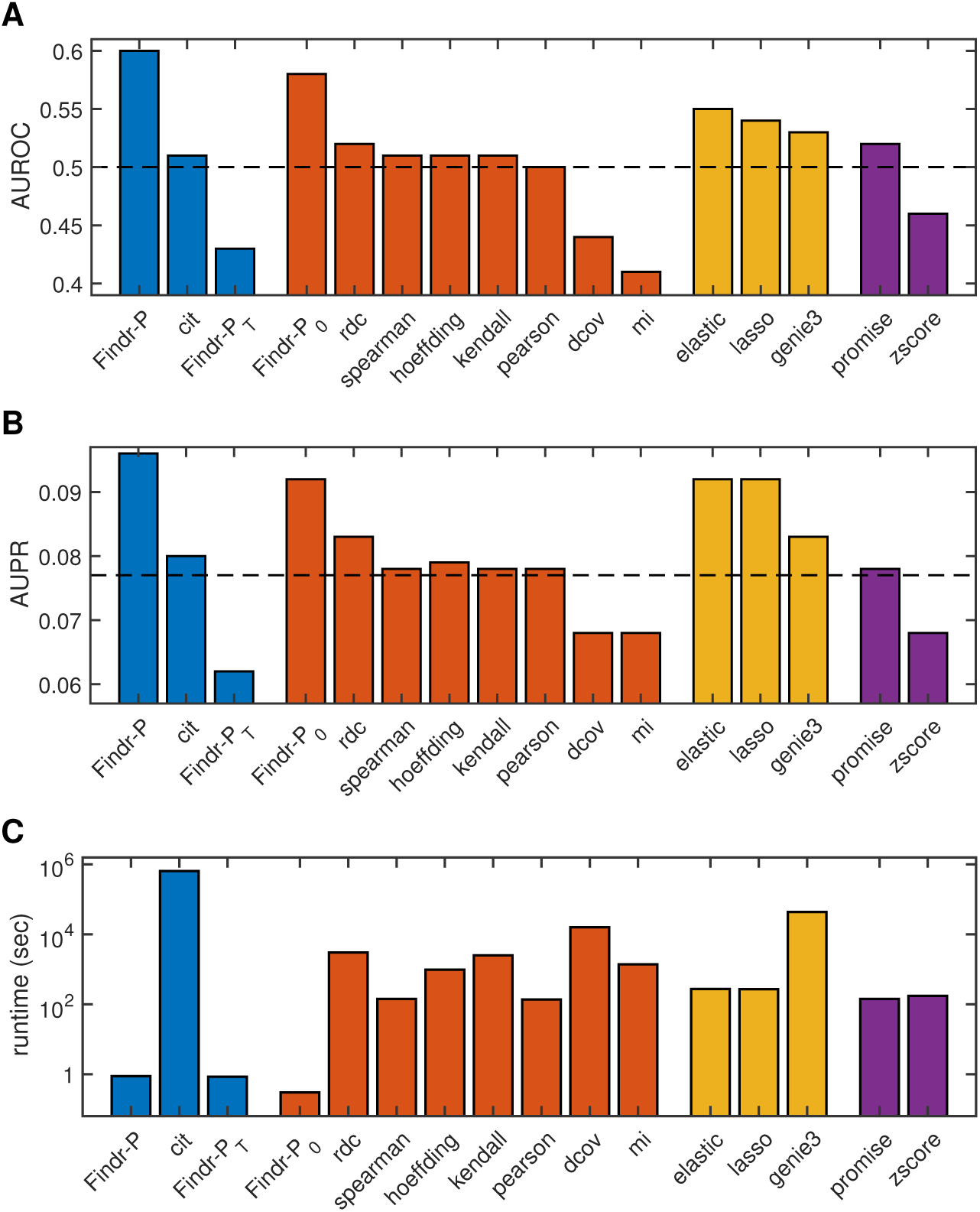
Findr achieves highest accuracy and speed on the prediction of miRNA target genes from the Geuvadis data. Shown are the AUROC (**A**), AUPR (**B**) and runtime (**C**) for 16 miRNA target prediction methods. Methods are colored by type: blue, genotype-assisted causal inference methods; red, pairwise correlation methods; yellow, multivariate regression methods; purple, other methods. Dashed lines are the AUROC and AUPR from random predictions. For method details, see **Supplementary Material S2.4**.

### 2.6 Findr predicts transcription factor targets with more accurate FDR estimates

We considered 3,172 genes with significant cis-eQTLs in the Geuvadis data^3^ (Section 4.1) and inferred regulatory interactions to the 23,722 target genes using Findr’s traditional (*P*_*T*_), new (*P*) and correlation (*P*_0_) tests, and CIT. Groundtruths of experimentally confirmed causal gene interactions in human, and mammalian systems more generally, are of limited availability and mainly concern transcription or transcription-associated DNA-binding factors (TFs). Here we focused on a set of 25 TFs in the set of eQTL-genes for which either differential expression data following siRNA silencing (6 TFs) or TF-binding data inferred from ChIP-sequencing and/or DNase foot-printing (20 TFs) in a lymphoblastoid cell line (GM12878) was available^25^ (Section 4.1). AUPRs and AUROCs did not exhibit substantial differences, other than modest improvement over random predictions (**Figure S9**). To test for enrichment of true positives among the top-ranked predictions, which would be missed by global evaluation measures such as AUPR or AUROC, we took advantage of the fact that Findr’s probabilities are empirical local precision estimates for each test (Section 4.5), and assessed how estimated local precisions of new, traditional, and correlation tests reflected the actual precision. Findr’s new test correctly reflected the precision values at various threshold levels, and was able to identify true regulations at high precision control levels (Figure 4). However, the traditional test significantly underestimated precision due to its elevated FNR. This lead to a lack of predictions at high precision thresholds but enrichment of true regulations at low thresholds, essentially nullifying the statistical meaning of its output probability *P*_*T*_. On the other hand, the correlation test significantly overestimated precisions because it is unable to distinguish causal, reversed causal or confounded interactions, which raises its FDR. The same results were observed when alternative groundtruth ChIP-sequencing networks were considered (**Figure S9**, **Figure S10**).

**Figure 4:**
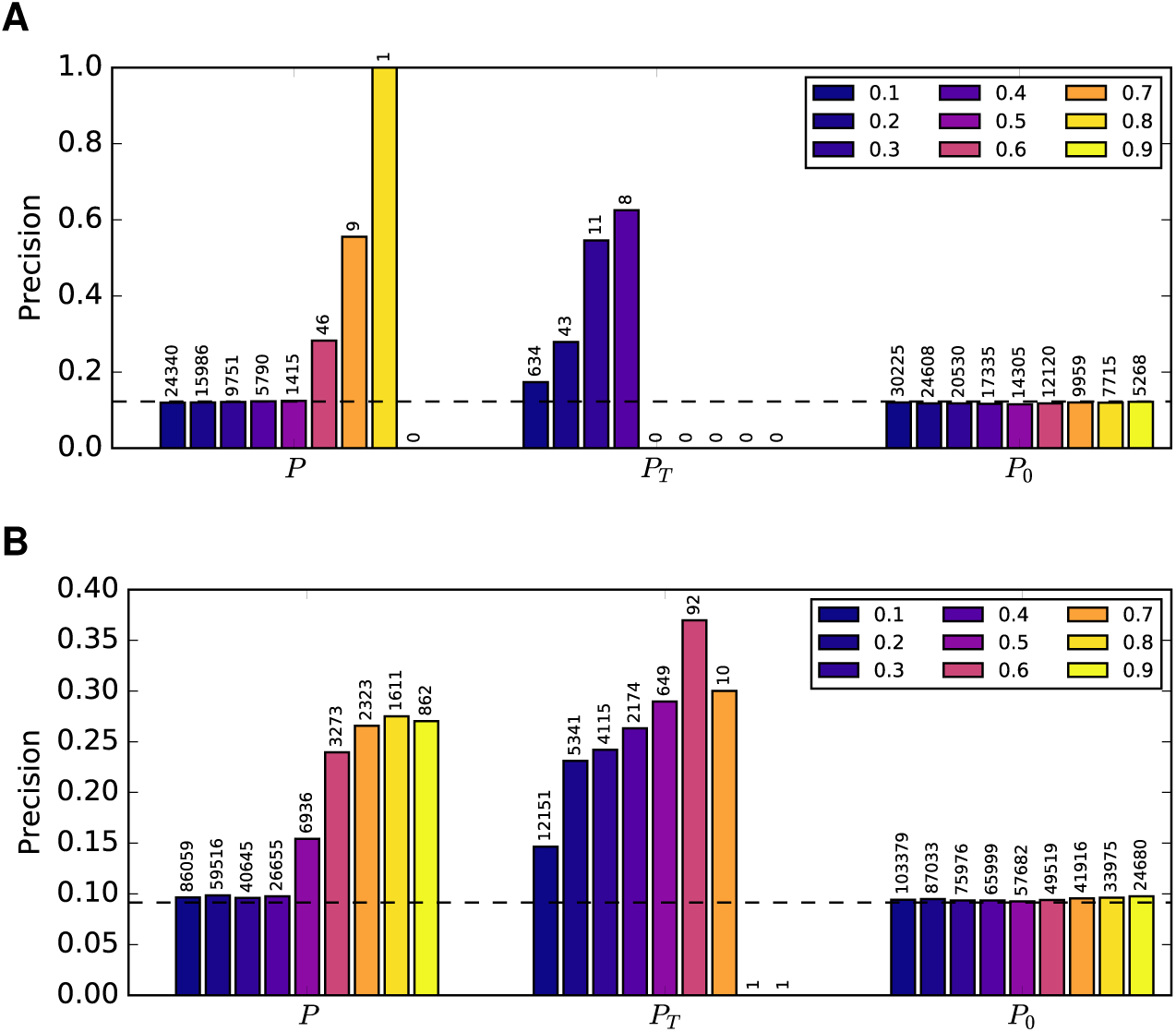
Findr predicts TF targets with more accurate FDR estimates from the Geuvadis data. The precision (i.e. 1-FDR) of TF target predictions is shown at probability cutoffs 0.1 to 0.9 (blue to yellow) with respect to known functional targets from siRNA silencing of 6 TFs (**A**) and known TF-binding targets of 20 TFs (**B**). The number above each bar indicates the number of predictions at the corresponding threshold. Dashed lines are precisions from random predictions.

## 3 Discussion

We developed a highly efficient, scalable software package Findr (Fast Inference of Networks from Directed Regulations) implementing novel and existing causal inference tests. Application of Findr on real and simulated genome and transcriptome variation data showed that our novel tests, which account for weak secondary linkage and hidden confounders at the potential cost of an increased number of false positives, resulted in a significantly improved performance to predict known gene regulatory interactions compared to existing methods, particularly traditional methods based on conditional independence tests, which had highly elevated false negative rates.

Causal inference using eQTLs as causal anchors relies on crucial assumptions which have been discussed in-depth elsewhere^8,9^. Firstly, it is assumed that genetic variation is always causal for variation in gene expression, or quantitative traits more generally, and is independent of any observed or hidden confounding factors. Although this assumption is valid for randomly sampled individuals, caution is required when this is not the case (e.g. case-control studies). Secondly, measurement error is assumed to be independent and comparable across variables. Correlated measurement error acts like a confounding variable, whereas a much larger measurement error in the source variable *A* than the target variable *B* may lead to an inversion of the inferred causal direction. The conditional independence test in particular relies on the unrealistic assumptions that hidden confounders and measurement errors are absent, the violation of which incurs false negatives and a failure to correctly predict causal relations, as shown throughout this paper.

Although the newly proposed test avoids the elevated FNR from the conditional independence test, it is not without its own limitations. Unlike the conditional independence test, the relevance and controlled tests (Table 1) are symmetric between the two genes considered. Therefore the direction of causality in the new test arises predominantly from using a different eQTL when testing the reverse interaction, potentially leading to a higher FDR as a minor trade-off. About 10% of cis-regulatory eQTLs are linked (as *cis*-eQTLs) to the expression of more than one gene^26^. In these cases, it appears that the shared *cis*-eQTL regulates the genes independently^26^, which in Findr is accounted for by the ‘controlled’ test (Table 1). When causality between genes and phenotypes or among phenotypes is tested, sharing or linkage of (e)QTLs can be more common. Resolving causality in these cases may require the use of Findr’s conservative, traditional causal inference test, in conjunction with the new test.

In this paper we have addressed the challenge of pairwise causal inference, but to reconstruct the actual pathways and networks that affect a phenotypic trait, two important limitations have to be considered. First, linear pathways propagate causality, and may thus appear as densely connected sets of triangles in pairwise causal networks. Secondly, most genes are regulated by multiple upstream factors, and hence some true edges may only have a small posterior probability unless they are considered in an appropriate multivariate context. The most straightforward way to address these issues would be to model the real directed interaction network as a Bayesian network with sparsity constraints. A major advantage of Findr is that it outputs probability values which can be directly incorporated as prior edge probabilities in existing network inference softwares.

In conclusion, Findr is a highly efficient and accurate open source software tool for causal inference from large-scale genome-transcriptome variation data. Its nonparametric nature ensures robust performances across datasets without parameter tuning, with easily interpretable output in the form of accurate precision and FDR estimates. Findr is able to predict causal interactions in the context of complex regulatory networks where unknown upstream regulators confound traditional conditional independence tests, and more generically in any context with discrete or continuous causal anchors.

## 4 Methods

### 4.1 Datasets

We used the following datasets/databases for evaluating causal inference methods:

1. Simulated genotype and transcriptome data of synthetic gene regulatory networks from the DREAM5 Systems Genetics challenge A (DREAM for short)^27^, generated by the SysGenSIM software^28^. DREAM provides 15 sub-datasets, obtained by simulating 100, 300, and 999 samples of 5 different networks each, containing 1000 genes in every sub-dataset but more regulations for sub-datasets with higher numbering. In every sub-dataset, each gene has exactly one matching genotype variable. 25% of the genotype variables are cis-expression Quantitative Trait Loci (eQTL), defined in DREAM as: their variation changes the expression level of the corresponding gene directly. The other 75% are trans-eQTLs, defined as: their variation affects the expression levels of only the *downstream targets* of the corresponding gene, but not the gene itself. Because the identities of cis-eQTLs are unknown, we calculated the P-values of genotype-gene expression associations with kruX^29^, and kept all genes with a P-value less than 1/750 to filter out genes without cis-eQTL. For the subsampling analysis (detailed in Section 4.7), we restricted the evaluation to the prediction of target genes from these cis-genes only, in line with the assumption that Findr and other causal inference methods require as input a list of genes whose expression is significantly associated with at least one cis-eQTL. For the full comparison of Findr to the DREAM leaderboard results, we predicted target genes for all genes, regardless of whether they had a cis-eQTL.
2. Genotype and transcriptome sequencing data on 465 human lymphoblastoid cell line samples from the Geuvadis project^3^ consisting of the following data products:

- Genotype data (ArrayExpress accession E-GEUV-1)^30^.
- Gene quantification data for 23722 genes from nonredundant unique samples and after quality control and normalization (ArrayExpress accession E-GEUV-1)^31^.
- Quantification data of miRNA, with the same standard as gene quantification data (ArrayExpress accession E-GEUV-2)^32^.
- Best eQTLs of mRNAs and miRNAs (ArrayExpress accessions E-GEUV-1 and E-GEUV-2)^33,34^. We restricted our analysis to 360 European samples which are shared by gene and miRNA quantifications. Excluding invalid eQTLs from the Geuvadis analysis, such as single-valued genotypes, 55 miRNA-eQTL pairs and 3172 gene-eQTL pairs were retained.
3. For validation of predicted miRNA-gene interactions, we extracted the “strong” ground-truth table from miRLAB^23,35^, which contains experimentally confirmed miRNA-gene regulations from the following databases: TarBase^36^, miRecords^37^, miRWalk^38^, and miRTarBase^39^. The intersection of the Geuvadis and ground-truth table contains 20 miRNAs and 1054 genes with 1217 confirmed regulations, which are considered for prediction validation. Interactions that are present in the ground-truth table are regarded as true while others as false.
4. For verification of predicted gene-gene interactions, we obtained differential expression data following siRNA silencing of 59 transcription-associated factors (TFs) and DNA-binding data of 201 TFs for 8872 genes in a reference lymphoblastoid cell line (GM12878) from^25^. Six siRNA-targeted TFs, 20 DNA-binding TFs, and 6,790 target genes without missing differential expression data intersected with the set of 3172 eQTL-genes and 23722 target genes in Geuvadis and were considered for validation. We reproduced the pipeline of^25^ with the criteria for true targets as having a False Discovery Rate (FDR) < 0.05 from R package *qvalue* for differential expression in siRNA silencing, or having at least 2 TF-binding peaks within 10kb of their transcription start site. We also obtained the filtered proximal TF-target network from^40^, which had 14 TFs and 7,000 target genes in common with the Geuvadis data.

### 4.2 General inference algorithm

Consider a set of observations sampled from a mixture distribution of a null and an alternative hypothesis. For instance in gene regulation, every observation can correspond to expression levels of a pair of genes wich are sampled from a bivariate normal distribution with zero (null hypothesis) or non-zero (alternative hypothesis) correlation coefficient. In Findr, we predict the probability that any sample follows the alternative hypothesis with the following algorithm (based on and modified from^11^):

1. For robustness against outliers, we convert every continuous variable into standard normally distributed *N*(0,1) values using a rank-based inverse normal transformation across all samples. We name this step as *supernormalization*.
2. We propose a null and an alternative hypothesis for every likelihood ratio test (LRT) of interest where, by definition, the null hypothesis space is a subset of the alternative hypothesis. Model parameters are replaced with their maximum likelihood estimators (MLEs) to obtain the log likelihood ratio (LLR) between the alternative and null hypotheses (Section 4.3).
3. We derive the analytical expression for the probablity density function (PDF) of the LLR when samples follow the null hypothesis (Section 4.4).
4. We convert LLRs into posterior probabilities of the hypothesis of interest with the empirical estimation of local FDR (Section 4.5).

Implementational details can be found in Findr’s source code.

### 4.3 Likelihood ratio tests

Consider correlated genes *A, B*, and a third variable *E* upstream of *A* and *B*, such as a significant eQTL of *A*. The eQTLs can be obtained either *de novo* using eQTL identification tools such as matrix-eQTL^41^ or kruX^29^, or from published analyses. Throughout this article, we assume that *E* is a significant eQTL of *A*, whereas extension to other data types is straightforward. We use *A*_*i*_ and *B*_*i*_ for the expression levels of gene *A* and *B* respectively, which are assumed to have gone through the supernormalization in Section 4.2, and optionally the genotypes of the best eQTL of *A* as *E_j_*, where *i* = 1,…,*n* across samples. Genotypes are assumed to have a total of *n*_*a*_ alleles, so *E*_*i*_ ∈ {0,…, *n*_*a*_}. We define the null and alternative hypotheses for a total of six tests, as shown in Table 1. LLRs of every test are calculated separately as follows:

0. **Correlation test:** Define the null hypothesis as *A* and *B* are independent, and the alternative hypothesis as they are correlated:

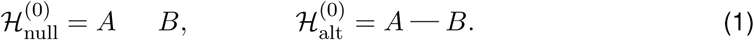

The superscript (0) is the numbering of the test. Both hypotheses are modeled with gene expression levels following bivariate normal distributions, as

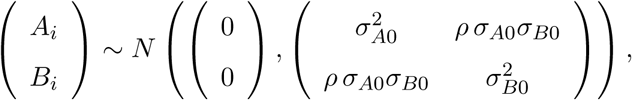
for *i* = 1,…, *n*. The null hypothesis corresponds to *ρ* = 0.

Maximum likelihood estimators (MLE) for the model parameters *ρ*, σ_*A*0_, and *B_Β0_* are

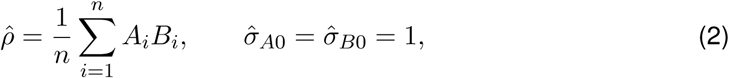
and the LLR is simply

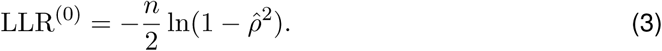

In the absence of genotype information, we use nonzero correlation between *A* and *B* as the indicator for *A* → *B* regulation, giving the posterior probability

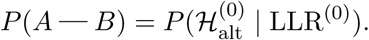
false negative

1. **Primary (linkage) test:** Verify that *E* regulates *A* from 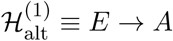 and 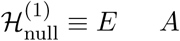. For 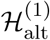, we model *E* → *A* as *A* follows a normal distribution whose mean is determined by *E* categorically, i.e.

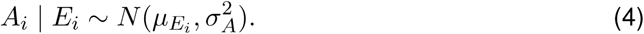 From the total likelihood 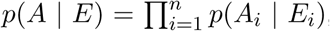, we find MLEs for model parameters *μ*_*j*_, j = 0, 1,…, *n*_*a*_, and σ_*A*_, as

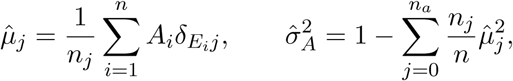
where *n*_*j*_ is the sample count by genotype category,

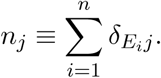 The Kronecker delta function is defined as *δ*_*xy*_ = 1 for *x* = *y*, and 0 otherwise. When summing over all genotype values (*j* = 0,…, *n*_*a*_), we only pick those that exist (*n*_*j*_ > 0) throughout this article. Since the null hypothesis is simply that *A*_*i*_ is sampled from a genotype-independent normal distribution, with MLEs of mean zero and standard deviation one due to the supernormalization (Section 4.2), the LLR for test 1 becomes

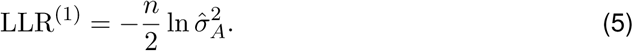 By favoring a large LLR^(1)^, we select 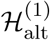 and verify that *E* regulates *A*, with

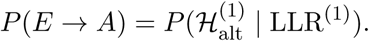
2. **Secondary (linkage) test:** The secondary test is identical with the primary test, except it verifies that *E* regulates *B*. Hence repeat the primary test on *E* and *B* and obtain the MLEs:

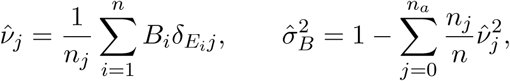
and the LLR as

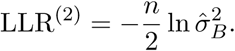 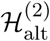 is chosen to verify that *E* regulates *B*.
3. **(Conditional) independence test:** Verify that *E* and *B* are independent when conditioning on *A*. This can be achieved by comparing 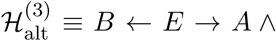 (*A* correlates with *B*) against 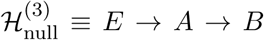. LLRs close to zero then prefer 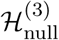, and ensure that *E* regulates *B* only through *A*:

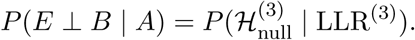 For 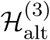, the bivariate normal distribution dependent on *E* can be represented as

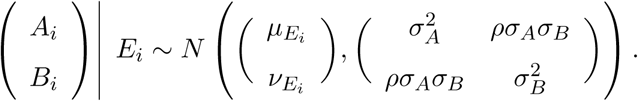 For 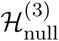, the distributions follow Eq 4, as well as

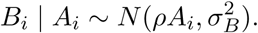 Substituting parameters μ_*j*_, *v*_*j*_, *σ*_*A*_, *σ*_*B*_, *ρ* of 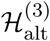 and μ_*j*_, *ρ, σ*_*A*_, *σ*_*B*_ of 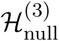 with their MLEs, we obtain the LLR:

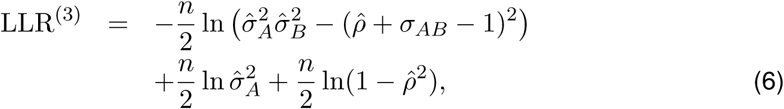
where

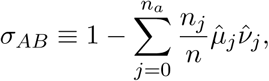
and 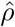 is defined in Eq 2.
4. **Relevance test**: Since the indirect regulation *E → B* tends to be weaker than any of its direct regulation components (*E* → *A* or *A* → *B*), we propose to test *E* → *A → B* with indirect regulation *E* → *B* as well as the direct regulation *A* → *B* for stronger distinguishing power on weak regulations. We define 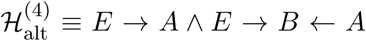 and 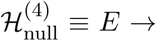 *A B*. This simply verifies that *B* is not independent from both *A* and *E* simultaneously. In the alternative hypothesis, *B* is regulated by *E* and *A*, which is modeled as a normal distribution whose mean is additively determined by *E* categorically and *A* linearly, i.e.

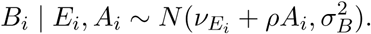 We can hence solve its LLR as

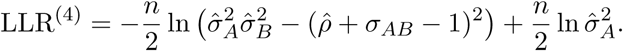
5. **Controlled test**: Based on the positives of the secondary test, we can further distinguish the alternative hypothesis 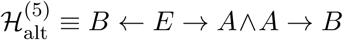 from the null 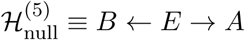 to verify that *E* does not regulate *A* and *B* independently. Its LLR can be solved as

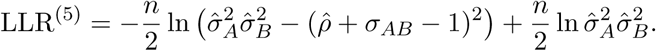

### 4.4 Null distributions for the log-likelihood ratios

The null distribution of LLR, 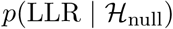, may be obtained either by simulation or analytically. Simulation, such as random permutations from real data or the generation of random data from statistics of real data, can deal with a much broader range of scenarios in which analytical expressions are unattainable. However, the drawbacks are obvious: simulation can take hundreds of times longer than analytical methods to reach a satisfiable precision. Here we obtained analytical expressions of 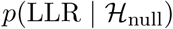 for all the tests introduced above.

0. **Correlation test**: 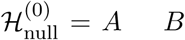 indicates no correlation between *A* and *B*. Therefore, we can start from

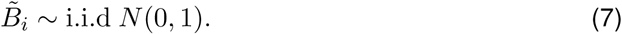

In order to simulate the supernormalization step, we normalize 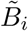 into *B_i_* with zero mean and unit variance as:

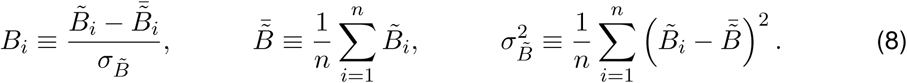

Transform the random variables 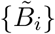 by defining

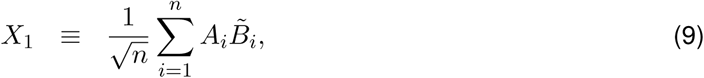

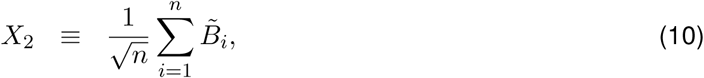

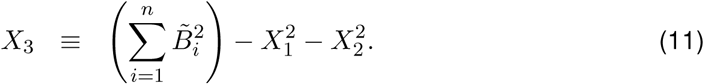

Since 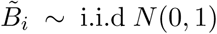 (according to Eq 7), we can easily verify that *X*_1_, *X*_2_, *X*_3_ are independent, and

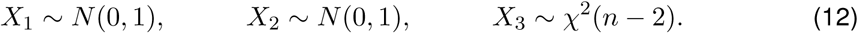

Expressing Eq 3 in terms of *X*_1_, *X*_2_, *X*_3_ gives

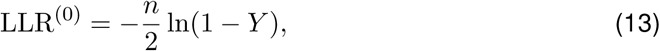
in which

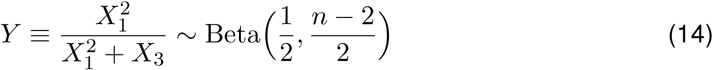
follows the Beta distribution.

We define distribution 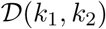 as the distribution of a random variable 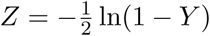 for *Y* ∼ Beta(*k*_1_/2, *k*_2_/2), i.e.

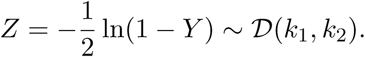

The probability density function (PDF) for 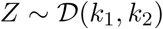 can be derived as: for *z* > 0,

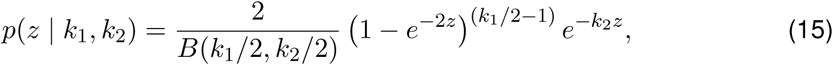
and for *z* ≤ 0, *p*(*z* | *k*_1_,*k*_2_) = 0. Here *B*(*a*,*b*) is the Beta function. Therefore the null distribution for the correlation test is simply

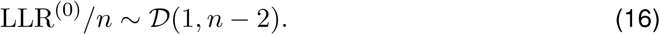

1. **Primary test**: 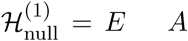 indicates no regulation from *E* to *A*. Therefore, similarly with the correlation test, we start from 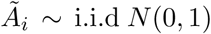 and normalize them to *A*_*i*_ with zero mean and unit variance.

The expression of LLR^(1)^ then becomes:

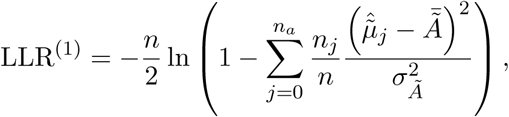

Where

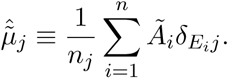

For now, assume all possible genotypes are present, i.e. *n*_*j*_ > 0 for *j* = 0,…, *n*_*a*_. Transform 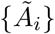 by defining

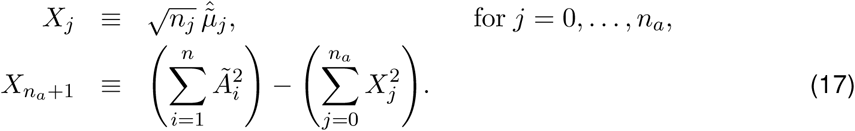

Then we can similarly verify that {*X*_*i*_} are pairwise independent, and

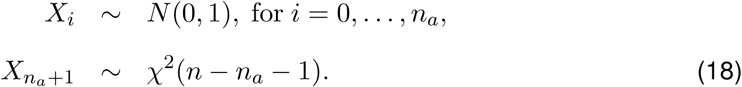

Again transform {*X*_*i*_} by defining independent random variables

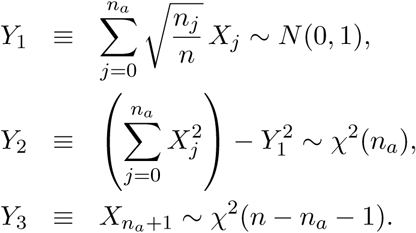

Some calculation would reveal

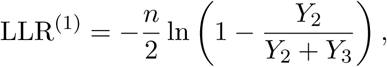

i.e.

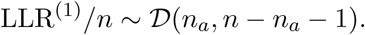

To account for genotypes that do not show up in the samples, define 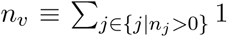 as the number of different genotype values across all samples. Then

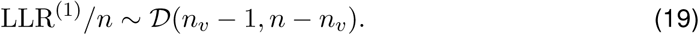

2. **Secondary test**: Since the null hypotheses and LLRs of primary and secondary tests are identical, LLR^(2)^ follows the same null distribution as Eq 19.

3. **Independence test**: The independence test verifies if *E* and *B* are uncorrelated when conditioning on *A*, with 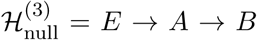. For this purpose, we keep *E* and *A* intact while randomizing 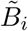 according to *B*’s correlation with *A*:

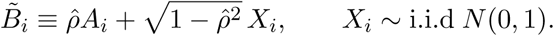

Then 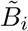 is normalized to *B*_*i*_ according to Eq 8. The null distribution of LLR^(3)^ can be obtained with similar but more complex computations from Eq 6, as

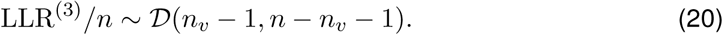

4. **Relevance test**: The null distribution of LLR^(4)^ can be obtained similarly by randomizing *B*_*i*_ according to Eq 7 and Eq 8, as

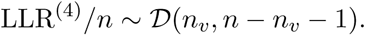

5. **Controlled test**: To compute the null distribution for the controlled test, we start from

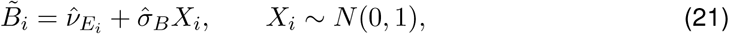
and then normalize 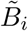 into *B*_*i*_ according to Eq 8. Some calculation reveals the null distribution as

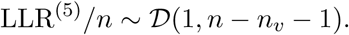

We verified our analytical method of deriving null distributions by comparing the analytical null distribution v.s. null distribution from permutation for the relevance test in **Section S2.2**.

### 4.5 Bayesian inference of posterior probabilities

After obtaining the PDFs for the LLRs from real data and the null hypotheses, we can convert LLR values into posterior probabilities 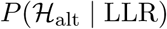. We use a similar technique as in^11^, which itself was based on a more general framework to estimate local FDRs in genome-wide studies^42^. This framework assumes that the real distribution of a certain test statistic forms a mixture distribution of null and alternative hypotheses. After estimating the null distribution, either analytically or by simulation, it can be compared against the real distribution to determine the proportion of null hypotheses, and consequently the posterior probability that the alternative hypothesis is true at any value of the statistic.

To be precise, consider an arbitrary likelihood ratio test. The fundamental assumption is that in the limit LLR → 0^+^, all test cases come from the null hypothesis 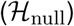, whilst as LLR increases, the proportion of alternative hypotheses 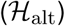 also grows. The mixture distribution of real LLR values is assumed to have a PDF as

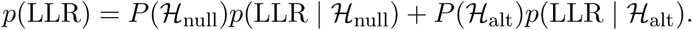

The priors 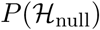 and 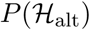 sum to unity and correspond to the proportions of null and alternative hypotheses in the mixture distribution. For any test *i* = 0,…, 5, Bayes’ theorem then yields its posterior probability as

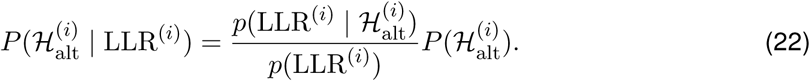

Based on this, we can define the posterior probabilities of the selected hypotheses according to Table 1, i.e. the alternative for tests 0, 1, 2, 4, 5 and the null for test 3 as

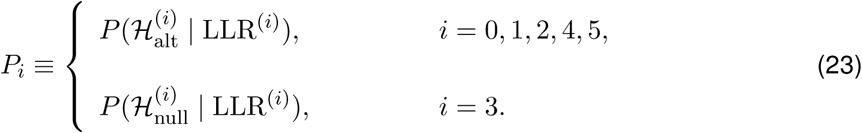

After obtaining the LLR distribution of the null hypothesis 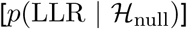, we can determine its proportion 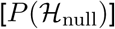 by aligning 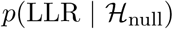 with the real distribution *p*(LLR) at the LLR → 0^+^ side. This provides all the prerequisites to perform Bayesian inference and obtain any *P_i_* from Eq 23.

In practice, PDFs are approximated with histograms. This requires proper choices of histogram bin widths, 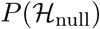, and techniques to ensure the conversion from LLR to posterior probability is monotonically increasing and smooth. Implementational details can be found in Findr package and in **Section S1.1.** Distributions can be estimated either separately for every (*E, A*) pair or by pooling across all (*E, A*) pairs. In practice, we test on the order of 10^3^ to 10^4^ candidate targets (“*B*”) for every (*E, A*) such that a separate conversion of LLR values to posterior probabilities is both feasible and recommended, as it accounts for different roles of every gene, especially hub genes, through different rates of alternative hypotheses.

Lastly, in a typical application of Findr, inputs of (*E, A*) pairs will have been pre-determined as the set of significant eQTL-gene pairs from a genome-wide eQTL associaton analysis. In such cases, we may naturally assume *P*_1_ = 1 for all considered pairs, and skip the primary test.

### 4.6 Tests to evaluate

Based on the six tests in Section 4.3, we use the following tests and test combinations for the inference of genetic regulations, and evalute them in the results.

- The correlation test is introduced as a benchmark, against which we can compare other methods involving genotype information. Pairwise correlation is a simple measure for the probability of two genes being functionally related either through direct or indirect regulation, or through coregulation by a third factor. Bayesian inference additionally considers different gene roles. Its predicted posterior probability for regulation is *P*_0_.
- The traditional causal inference test, as explained in^11^, suggested that the regulatory relation *E* → *A* → *B* can be confirmed with the combination of three separate tests: *E* regulates *A, E* regulates *B*, and *E* only regulates *B* through *A* (i.e. *E* and *B* become independent when conditioning on *A*). They correspond to the primary, secondary, and independence tests respectively. The regulatory relation *E* → *A* → *B* is regarded positive only when all three tests return positive. The three tests filter the initial hypothesis space of all possible relations between *E, A*, and *B*, sequentially to *E* → *A* (primary test), *E* → *A* Λ *E* → *B* (secondary test), and *E* → *A* → *B* Λ (no confounder for *A* and B) (conditional independence test). The resulting test is stronger than *E* → *A* → *B* by disallowing confounders for *A* and *B*. So its probability can be broken down as

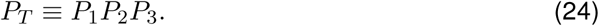 Trigger^43^ is an R package implementation of the method. However, since Trigger integrates eQTL discovery with causal inference, it is not practical for use on modern datasets. For this reason, we reimplemented this method in Findr, and evaluated it with *P*_2_ and *P*_2_*P*_3_ separately, in order to assess the individual effects of secondary and independence tests. As discussed above, we expect a set of significant eQTLs and their associated genes as input, and therefore *P*_1_ = 1 is assured and not calculated in this paper or the package Findr. Note that *P*_*T*_ is the estimated local precision, i.e. the probability that tests 2 and 3 are both true. Correspondinly, its local FDR (the probability that one of them is false) is 1 − *P*_*T*_.
- The novel test, aimed specifically at addressing the failures of the traditional causal inference test, combines the tests differently:

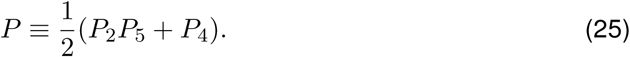

Specifically, the first term in Eq 25 accounts for hidden confounders. The controlled test replaces the conditional independence test and constrains the hypothesis space more weakly, only requiring the correlation between *A* and *B* is not entirely due to pleiotropy. Therefore, *P*_2_*P*_5_ (with *P*_1_ = 1) verifies the hypothesis that 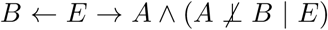, a superset of *E* → *A* → *B*.

On the other hand, the relevance test in the second term of Eq 25 addresses weak interactions that are undetectable by the secondary test from existing data (*P*_2_ close to 0). This term still grants higher-than-null significance to weak interactions, and verifies that *E* → *A* Λ (*E* → *B* V *A* — *B*), also a superset of *E* → *A* → *B*. In the extreme undetectable limit where *P*_2_ = 0 but *P*_4_ ≠ 0, the novel test Eq 25 automatically reduces to 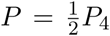, which assumes equal probability of either direction and assigns half of the relevance test probability to *A* → *B*.

The composite design of the novel test aims not to miss any genuine regulation whilst distinguishing the full spectrum of possible interactions. When the signal level is too weak for tests 2 and 5, we expect *P*_4_ to still provide distinguishing power better than random predictions. When the interaction is strong, *P*_2_*P*_5_ is then able to pick up true targets regardless of the existence of hidden confounders.

### 4.7 Evaluation methods

- **Evaluation metrics:** Given the predicted posterior probabilities for every pair (*A, B*) from any test, or more generically a score from any inference method, we evaluated the predictions against the direct regulations in the ground-truth tables (Section 4.1) with the metrics of Receiver Operating Characteristic (ROC) and Precision-Recall (PR) curves, as well as the Areas Under the ROC (AUROC) and Precision-Recall (AUPR) curves^44^. In particular, AUPR is calculated with the Davis-Goadrich nonlinear interpolation^45^ with R package *PRROC*.
- **Subsampling:** In order to assess the effect of sample size on the performances of inference methods, we performed subsampling evaluations. This is made practically possible by the DREAM datasets which contain 999 samples with sufficient variance, as well as the computational efficiency from Findr which makes subsampling computationally feasible. With a given dataset and ground-truth table, the total number of samples n, and the number of samples of our actual interest *N* < *n*, we performed subsampling by repeating following steps k times:
  1. Randomly select *N* samples out of the total *n* samples without replacement.
  2. Infer regulations only based on the selected samples.
  3. Compute and record the evaluation metrics of interest (e.g. AUROC and AUPR) with the inference results and ground-truths.
  Evaluation metrics are recorded in every loop, and their means, standard deviations, and standard errors over the *k* runs, are calculated. The mean indicates how the inference method performs on the metric in average, while the standard deviation reflects how every individual subsampling deviates from the average performance.
- **Local precision of top predictions separately for confounded and unconfounded gene pairs:** In order to demonstrate the inferential precision among top predictions for any inference test (here the traditional and novel tests separately), we first ranked all (ordered) gene pairs (*A, B*) according to the inferred significance for *A* → *B*. All gene pairs were split into groups according to their relative significance ranking (9 groups in Figure 1C,D, as top 0% to 0.01%, 0.01% to 0.02%, etc). Each group was divided into two subgroups, based on whether each gene pair shared at least one direct upstream regulator gene (confounded) or not (unconfounded), according to the gold standard. Within each subgroup, the local precision was computed as the number of true directed regulations divided by the total number of gene pairs in the subgroup.

### 4.8 Simulation studies on causal models with measurement error

We investigated how each statistical test tolerates measurement errors with simulations in a controlled setting. We modelled the causal relation *A* → *B* in a realistic setup as *E* → *A*^(t)^ → *B* with *A*^(t)^ → *A*. *E* remains as the accurately measured genotype values as the eQTL for the primary target gene *A*. *A*^(t)^ is the true expression level of gene *A*, which is not observable. *A* is the measured expression level for gene *A*, containing measurement errors. *B* is the measured expression level for gene *B*.

For simplicity, we only considered monoallelic species. Therefore the genotype *E* in each sample followed the Bernoulli distribution, parameterized by the predetermined minor allele frequency. Each regulatory relation (of *E* → *A*^(t)^, *A*^(t)^ → *A*, and *A*^(t)^ → *B*) correponded to a normal distribution whose mean was linearly dependent on the regulator variable. In particular, for sample *i*:

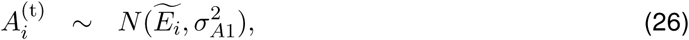

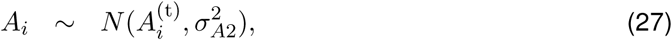

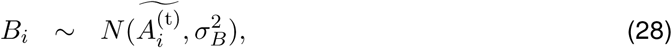
in which σ_*A*1_, σ_*A*2_, and σ_*B*_ are parameters of the model. Note that 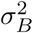 is *B*’s variance from all unknown sources, including expression level variations and measurement errors. The tilde normalizes the variable into zero mean and unit variance, as:

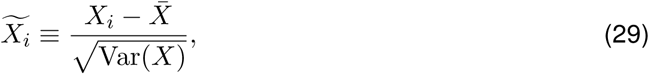
where 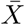 and Var(*X*) are the mean and variance of *X* ≡ {*X*_*i*_} respectively.

Given the five parameters of the model (the number of samples, the minor allele frequency, σ_*A*1_, σ_*A*2_, and σ_*B*_), we could simulate the observed data for *E, A*, and *B*, which were then fed into Findr for tests 2-5 and their p-values of the respective null hypotheses. Supernormalization step was replaced with normalization which merely shifted and scaled variables into zero mean and unit variance.

We then chose different configurations on the number of samples, the minor allele frequency, and σ_*B*_. For each configuration, we varied σ_*A*1_ and σ_*A*2_ in a wide range to obtain a 2-dimensional heatmap plot for the p-value of each test, thereby exploring how each test was affected by measurement errors of different strengths. Only tiles with a significant *E* → *A* eQTL relation were retained. The same initial random seed was employed for different configurations to allow for replicability.

## Acknowledgements

This work was supported by the BBSRC (grant numbers BB/J004235/1 and BB/M020053/1).

